# All-in-one comprehensive extraction of metabolites, proteins and ribonucleic acids for the rapid analysis of oil palm systems biology

**DOI:** 10.1101/2020.07.01.183475

**Authors:** Syahanim Shahwan, Abrizah Othman, Zain Nurazah, Nurul Liyana Rozali, Umi Salamah Ramli

## Abstract

Oil palm (*Elaeis guineensis* Jacq.) systems biology offers a comprehensive view of the plant system by employing a holistic multi-omics approach encompassing the molecular data at various hierarchical levels. Sample limitation and the importance of integrating all molecular data with minimal variation, led to the development of sequential extraction of biomolecule fractions from a single undivided biological sample. This article describes a workflow for the comprehensive isolation of metabolites, proteins and ribonucleic acids from oil palm root. Samples were subjected to solvent extraction with methanol-chloroform-water to recover metabolites of diverse polarity. The resultant pellet was subjected to buffer and solvent partitioning to obtain RNA and proteins. RNA extracted from the oil palm root showed a recovery of 180.25 ng mg^-1^, with a A260:A280 ratio ranging between1.9-2.0 and a RIN value of 6.7. Co-extracted proteins resulted in a recovery of 29.28 μg mg^-1^ and revealed a total of 1852 identified proteins. Polar metabolites revealed approximately 40 metabolite peaks, and non-polar metabolites with two major fatty acid groups i.e. saturated and unsaturated fatty acids at 55.4% and 38.6%, respectively. This protocol demonstrated an advancement of extraction protocols for oil palm root biomolecules, which will consecutively expedite the establishment of various multi-omics platforms.

**Highlight:** Metabolites, proteins and RNA are co-extracted from oil palm root using the all-in-one extraction protocol which provides biomolecule extracts for various omics platforms.

## Introduction

A comprehensive study and analysis of complex biological components within tissues or cells, provides a holistic approach to comprehend complex cellular processes and functions (Bruggeman and Westerhoff, 2007). Biological input obtained from the various omics platforms including transcriptomics, proteomics and metabolomics are integrated to understand the interrelationship between networks of biological processes (Potters, 2010). A new improved breeding material is always in demand by oil palm stakeholders. Thus, improving oil palm varieties via its genetics information is crucial and need to be materialized by advanced biotechnology (Kushairi *et al*., 2019). Omics-based studies have been fully endeavored in oil palm research with extensive discoveries of transcripts (Low *et al*., 2008; Singh *et al*., 2013; Tee *et al*., 2013; Singh *et al*., 2014; Xia *et al*., 2014; Ooi *et al*., 2015; Bahari *et al*., 2018; Rosli *et al*., 2018; Avila-Mendez *et al*., 2019), proteins (Syahanim *et al*., 2013; Jeffery Daim *et al*., 2015; Ooi *et al*., 2015; Lau *et al*., 2018; Hassan *et al*., 2019) and metabolites (Tahir *et al*., 2012; Zain *et al*., 2013; Rozali *et al*., 2017). It is imperative to understand the molecular mechanisms governing the complex diversity of biomolecules and dynamics in oil palm traits such as yield, height, quality and disease resistance using different oil palm tissues including leaf, root, fruit, seed, trunk and basal stem, which can be deployed in breeding programmes. Transcriptomic analysis has been widely used in exploring the expression of mRNA in oil palm clonal materials under various biotic and abiotic stress conditions. As reported by Avila-Mendez *et al*. (2019), transcriptional analysis has provided an overview of the genes involved in the molecular response of the oil palm clones to *Phytophra palmivora*. Shearman *et al*. (2019) analysed expression data for all transcripts in flowers and fruits of mantled and normal oil palm clones in order to identify differentially expressed genes between these two populations.

Integration of knowledge driven by omics-based datasets has been initiated in order to understand biological system/responses under favourable or unfavourable conditions. Two or more omics datasets have been utilised to dissect strong and novel biomolecule candidates associated with the molecular process of interest. Pairwise analysis of metabolite and transcript datasets of potato tuber systems revealed significant correlation and included several strong correlations to novel nutritionally important metabolites (Urbanczyk-Wochniak *et al*., 2003). Gene-to-gene interaction and metabolite-to-gene networking was elucidated in *Arabidopsis thaliana* grown under sulphur deficiency, via integrated metabolomics and transcriptomics (Hirai *et al*., 2005). Omics-based datasets have also been used in medicinal plant research and revealed novel biomolecules which were further functionally characterised (Ziegler *et al*., 2005; Rischer *et al*., 2006; Ziegler *et al*., 2006; Gesell *et al*., 2009; Liscombe *et al*., 2009; Pienkny *et al*., 2009; Hagel and Facchini, 2013; Saito, 2013; Srivastava *et al*., 2013; Chen and Facchini, 2014; Marques *et al*., 2014; Miettinen *et al*., 2014; Rohani *et al*., 2016; Udomsom *et al*., 2016; Rai *et al*., 2017). Integration of omics-based datasets were also used for dissecting biological response of *Brassicaceae* towards UV-B irradiation (Tohge *et al*., 2016). Multi-omics datasets of DNA methylome, transcriptome and metabolome were integrated and provided novel insight into the regulation of cotton fibre development by epigenetic mechanisms (Wang *et al*., 2016). Metabolite-based genome-wide-association-study (mGWAS), has been used to dissect the genetic control of plant metabolism in plants (Luo, 2015; Matsuda *et al*., 2015).

Analysis of biomolecules to understand the biochemical reactions within a cell involving transcripts, proteins and metabolites in oil palm systems biology is limited. Vidal (2009) indicated that the interaction of macromolecules and metabolites in cells or organs of an organism form a multi-scale dynamic complex system that is fundamental to biological processes. An integrated approach was developed in order to comprehend oil palm biomolecules as well as the compendium function of biological elements that are part of oil palm systems biology. At present, most of the integrative experiments involve the split-sample-study design where biological samples are divided into multiple parts with each part generating an omics dataset (Rai *et al*., 2017). Integration of the data is done later by using dedicated bioinformatics software tools to ascertain the desired interactions and functions in systems biology. But, this approach has limitations as it may introduce bias that leads to misinterpretation and miscorrelation as different populations of RNA, proteins and metabolites that might be expressed differently in the different sets of tissues (McConnell and Barton, 1998; Li *et al*., 2010; Eckert *et al*., 2012). Rai *et al*. (2017) iterated that combining multi-omics datasets poses great obstacles due to the requirements/criteria for scaling, noise removal, sensitivity and resolution for each data type variable. The authors also explained that different omics outputs result in lists of unrelated entities, and appropriate statistical tools need to be carefully chosen, in establishing biologically relevant relationships between cellular components. Experimental design is also a pivotal component, which significantly impacts the quality of omics-based datasets (Cavill *et al*., 2016). Thus, an integrated extraction protocol was designed in this study to extract RNA, proteins and metabolites in parallel. This protocol was also applied to overcome the major drawback of inadequate sample volume to perform various experiments via omics platforms. Chomczynski (1993) described the simultaneous extraction of RNA, DNA and proteins from human mammary epithelial cells, rat mammary tissues and mouse liver. Later, similar protocols were also employed to extract biomolecules from leaves and roots of plant tissues. For instance, a novel method for successful integrated extraction of metabolites, proteins and RNA from *Arabidopsis thaliana* was reported by Weckwerth *et al*. (2004). A universal protocol for the multiple molecular extraction of biomolecules (metabolites, DNA, total RNA, large RNA, small RNA, and proteins) from a single sample of a wide range of species such as *Populas, Pines, Arabidopsis* and *Chlamydomonas* was also described, allowing for a broad range of transcriptional studies, metabolite profiling and shotgun proteomics (Valledor *et al*., 2014). In addition, simultaneous extraction of biomolecules was employed in cell lines for a comprehensive molecular analysis of biological systems (Sapcariu *et al*., 2014; Vorreiter *et al*., 2016). A sequential isolation of metabolites, RNA, DNA and proteins applicable to microbial ecology was also presented to facilitate systematic multi-omics analysis and enable meaningful data integration (Roume *et al*., 2013).

This report describes an all-in-one extraction protocol of metabolites, proteins and RNA from a single oil palm root sample. The developed protocol could then be applied to understanding the oil palm systems biology through the integration of multi-omics data: transcriptomics, proteomics and metabolomics simultaneously, thus minimizing the variance that would result from multiple experiments.

## Materials and Methods

### Plant materials

Oil palm (*Deli dura* X *Avros pisifera*) root samples were obtained from MPOB Kluang Research Station, Johor, Malaysia in three biological replicates. The roots were cleared of soil and washed briefly. Another washing step was performed using sterile water, and primary roots were cut into small pieces and then snap frozen using liquid nitrogen and stored at −80°C until use.

### All-in-one-extraction: At a glance

Fig. 1 represents a schematic overview of the experimental workflow of the established protocol which can be easily adjusted in accordance to the required quantity of samples. At a glance, solvent extraction using a mixture of methanol, chloroform and water was used to extract metabolites of various range of polarities. The resulting pellet was washed and subjected to buffer/phenol/chloroform-based extraction for proteins and RNA. Later, buffer and phenol phases were precipitated to obtain the oil palm root RNA and protein, respectively. Qualitative analysis of the extracted biomolecules was evaluated using common and established analysis.

**Fig. 1.**
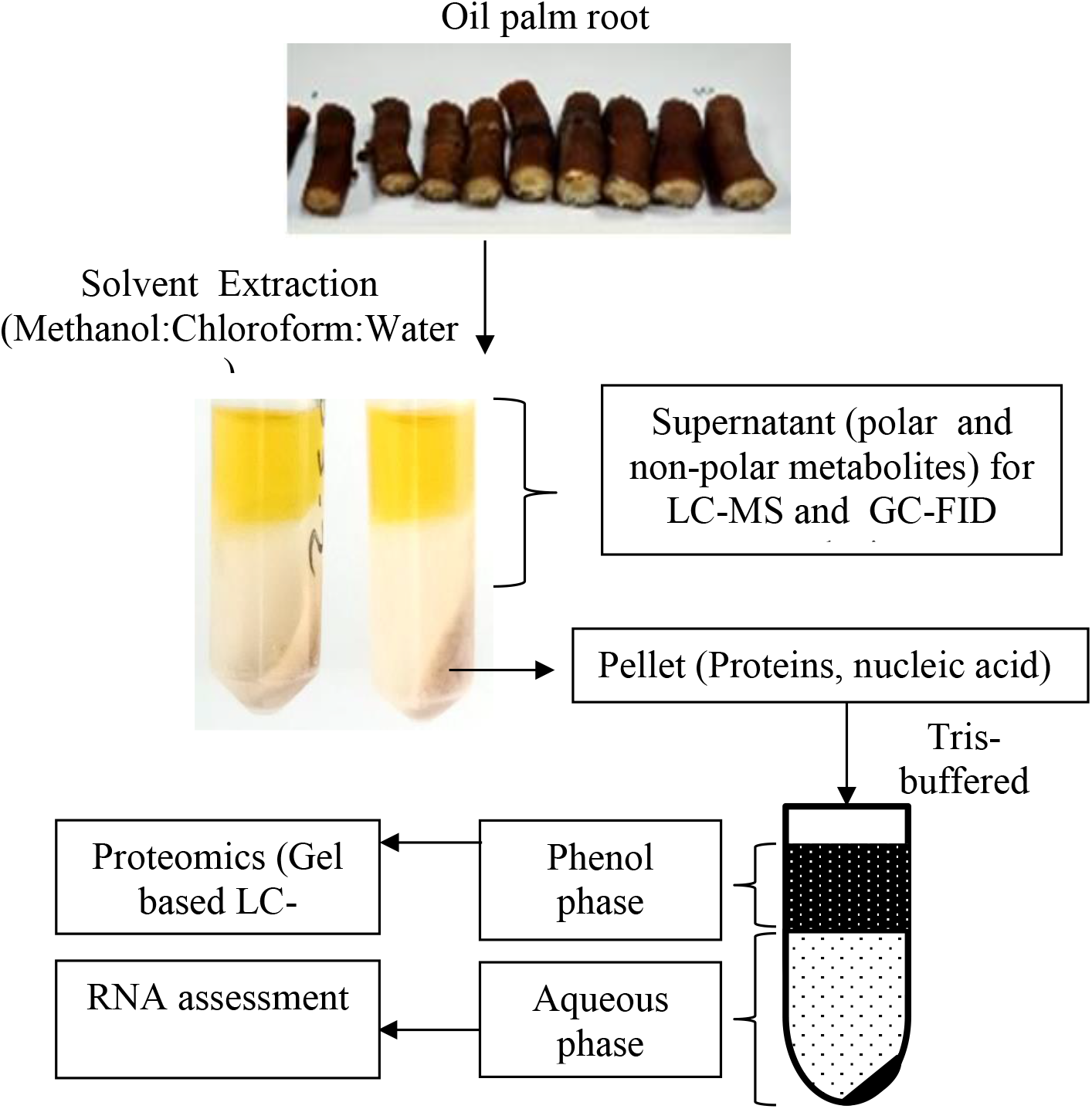
Schematic presentation of all-in-one extraction protocol of metabolites, proteins and RNA from the oil palm root.

### All-in-one-extraction: Metabolites

The integrated extraction protocol was developed for oil palm based on Weckwerth *et al*. (2004) and Valledor *et al*. (2014) with minor modification. Oil palm root (100 mg) was ground into a homogenous fine powder using mortar and pestle in liquid nitrogen. After adding a fixed volume (2.0 ml) of pre-cooled (20°C) solvent mixture of methanol, chloroform and water (2.5: 1:1), the tissues were thoroughly vortexed for 30 min and then centrifuged at 15000 *g* for 5 min. The resulting pellet was washed with 2.0 ml of pre-cooled chloroform/methanol mix (1:1). The solvent extracts were combined and used for metabolite analysis accordingly. Polar and non-polar components were separated by adding 5.0 ml of distilled water.

### All-in-one-extraction: RNA and Proteins

After removing the solvent phase from the tube, the pellet which consisted of nucleic acid (DNA/RNA), proteins, starch, membrane and cell wall components was resuspended with 1.0 ml protein extraction buffer. We tested three different buffers based on previous reports in order to identify the most suitable protein extraction buffer which gave reproducible results and minimised protein losses. The methods employed were adopted from Valledor *et al*. (2014), Weckwerth *et al*. (2004) and the modified method of Syahanim *et al*. (2013), and the buffers designated as *Buffer A, B* and *C*, respectively. The protein extraction buffer, i) *Buffer A* consisted of 0.1 M Tris-hydrochloride (Tris-HCl) pH 8.8, 25 mM ethylenediaminetetraacetic acid (EDTA), 0.9 M Sucrose and 0.4% (w/v) mercaptoethanol, ii) *Buffer B* consisted of 0.1 M Tris, 25 mM EDTA, 0.9 M sucrose and 0.4% (w/v) dithiothreitol (DTT) and iii) *Buffer C* consisted of 0.05 M Tris, 0.5 M sodium chloride (NaCl), 0.5% (v/v)-sodium dodecyl sulphate (SDS), 0.9 M sucrose and 0.4% (w/v) mercaptoethanol. Protease inhibitor was added to all buffer extracts to inhibit protease activity. The buffer mix was sonicated for 10 min at 4°C and added with 2.0 ml of Tris-buffered phenol. Then, the mix was vortexed for 10 min and centrifuged at 16 000 *g* for 10 min, portioned to two phases i.e. phenol phase and aqueous phase.

For ribonucleic acid extraction, 0.2 ml of chloroform was added to the aqueous phase. The mixtures were centrifuged at 16 000 *g* for 10 min at 4°C. Another 0.2 ml was added to the RNA-upper buffer phase and mixed briefly. After centrifugation at 16 000 *g* for 5 min, a total of 0.04 ml of acetic acid and 1.0 ml of ethanol were added to the aqueous phase for RNA precipitation overnight at −20°C. After centrifugation, the pellet was washed with 0.2 ml of 3 M sodium acetate and two times with 0.2 ml of 70% ethanol. The remaining pellet was re-dissolved in 50 μl RNase-free water.

The phenol phase which contained proteins was separated from the RNA-aqueous phase followed by back-extraction with an equal volume of protein extraction buffer. The phenol phase was precipitated with 5 volumes of pre-cooled 0.1 M ammonium acetate in 100% (v/v) methanol. The mixture was kept at −20°C for 16 hours and centrifuged at 12 000 *g* for 30 min at 4°C. The pellet was rinsed twice in 0.2 ml of methanol and twice with pre-cooled acetone. The protein pellet was dried for 5 min and dissolved in 0.5 to 1.0 ml solubilisation buffer containing 7 M Urea and 0.02 M DTT. Protein quantification was performed using Qubit Protein Assay Kit (Thermo Fisher Scientific, Wilmington, DE, USA) with bovine serum albumin (BSA) as standard at concentration range from 0 mg ml^-1^ to 5 mg ml^-1^.

### RNA integrity assessment

Total RNA was prepared according to the Agilent RNA 6000 Nano quick start guide and analysed using the Agilent BioAnalyzer 2100 (Agilent Technologies Inc). The RNA integrity number (RIN) was generated by the 2100 Software version B.02.08.SI648.

### Protein analysis by gel-based liquid chromatography-tandem mass spectrometry (LC-MS/MS)

Gel based LC MS was performed on protein samples using 12% polyacrylamide gel via Bio-Rad Mini-Protean III equipment (Bio-Rad) at 50 V for 15 min followed by 200 V until 1 cm mark below the stacking gel as described by Valledor and Weckwerth (2014b). Protein gels were stained for 30 min with Coomassie Brilliant Blue and destained 4 times for 80 min with destaining solution. Each lane was cut into two fragments and chopped into small pieces of 1 mm size. Blue colour was removed by soaking the gel pieces in 1 ml of 25 mM ammonium bicarbonate (NH_4_HCO_3_) for 15 min at 37°C and replaced with 1 ml of 25 mM NH_4_HCO_3_ in 50% (v/v) acetonitrile (ACN). The gel pieces were incubated at 37°C for 15 min. Solvent was discarded and the gel pieces were dehydrated with 200 μl of ACN for 5 min at room temperature. The gel pieces were dried using a SpeedVac followed by tryptic digestion at 2.5 ng μl^-1^ in trypsin buffer consisted of 25 mM NH_4_HCO_3_, 10% ACN and 5 mM calcium chloride (CaCl_2_). The peptide was digested at 37°C for 16 hours. The peptides were further extracted in three consecutive steps using 50% (v/v) ACN/1% (v/v) formic acid for two times and 90% (v/v) ACN/1% (v/v) formic acid. Supernatant was pooled and dried in the SpeedVac for 5 min. Peptide desalting was performed using a C18-ZipTip (Millipore, Bedford, MA) according to manufacturer’s recommendation.

The peptide was analysed using EASY-nLC 1000 (Thermo Fisher Scientific, Wilmington, DE, USA) coupled with Orbitrap Fusion mass spectrometry (Thermo Scientific, Wilmington, DE, USA). Chromatographic separations were performed on a reversed-phase analytical column (EASY-Spray Column Acclaim PepMap™ C18 100 A^0^, 2 μm particle size) (Thermo Scientific, Wilmington, DE, USA). Peptide separation was achieved *via* gradient settings at 5% to 40% solvent B (0.1% formic acid in ACN) for 91 min. The peptides were loaded onto the column with 95% solvent A (0.1% formic acid in LC-MS grade water) and desalting was performed for 1 min with 95% solvent A using reverse phase C18 trapping column. Tandem mass spectra were generated using the Orbitrap Fusion mass spectrometry (Thermo Scientific, Wilmington, DE, USA) and protein identification was performed using the Thermo Scientific™ Proteome Discoverer™ Software Version 2.1 against the oil palm in-house database and oil palm dataset available at UniProt. Database search parameter included the following: up to two missed cleavage sites was allowed; variable modification of oxidation (M), deamidation of asparagine (N) and glutamine (Q) and fixed modification, carbamidomethylation (C); peptide mass tolerance was 10 ppm. All peptides were validated using the percolator^®^ algorithm, based on q-value less than 1% false discovery rate (FDR).

### Polar metabolite analysis by liquid chromatography-mass spectrometry (LC-MS)

The polar phase containing polar metabolites was subjected to liquid chromatography-mass spectrometry (LC-MS) analysis. One microliter of the polar phase was injected to an Ultimate 3000 HPLC System coupled to MicrOTOF-Q™, a quadrupole/time of flight (Q/TOF) mass spectrometer (Bruker Daltonics, GmbH, Bremen, Germany). Separation was obtained on a Reverse-Phase Acclaim Polar Advantage II column (C18, 4.6 × 250 mm length, 5 μm particle size) (Thermo Scientific, Wilmington, DE, USA) at 37°C with a gradient elution programme of 1 ml min^-1^ flow rate with an increasing ACN/acetic acid gradient. All samples and replicates were continuously injected as one batch in random order to discriminate between technical and biological variations. Additionally, pooled samples were used as quality control (QCs) and injected at regular intervals throughout the analytical run. The Q/TOF MS analysis was performed in negative electrospray ionization and controlled by the HyStar Application version 3.2 software (Bruker Daltonics, GmbH, Bremen, Germany). The column effluent was set at 1.0 ml min^-1^. A split ratio of 1:4 was used generating a final flow rate of 250 μl min^-1^. Nitrogen was used as nebulising gas at 4.1 bar and 9.0 l min^-1^ flow rate. The temperature and voltage of capillary were set at 200°C and +3.5 kV, respectively. The full MS scan covered the mass range of *m/z* 50-1000.

The LC-MS data were processed through the Data Analysis 4.2 software (Bruker Daltonics, GmbH, Bremen, Germany). Find Molecular Features was applied to the raw data under these parameters: signal to noise ratio (S/N) threshold was set to 5.0, correlation coefficient was set to 0.7, minimum compound length was set to 10 spectra and smoothing width was set to 1,0. The data was evaluated in a time range from 0 to 40 min and in a mass range from *m/z* 50 to 1000.

### Non-polar metabolite analysis by gas chromatography-flame ionization detector (GC-FID)

The non-polar phase containing non-polar metabolites which included fatty acids was subjected to gas chromatography-flame ionization detection (GC-FID) (Clarus 500, Perkin Elmer). The non-polar phase was dried for fatty acid methyl esterification based on Yuan (2016). A total of 2 ml of 2.5% (v/v) sulfuric acid in methanol and 0.5 ml of toluene were dispensed to the extract prior to heating at 80°C for an hour. Then 2 ml of 0.9% NaCl and 1 ml of hexane were added and mixed together. The upper phase was transferred into a new vial and dried under nitrogen blower. Dried fatty acid methyl ester of oil palm root tissue was dissolved in 100 μl of hexane. One microliter of the derivatised fatty acid was then analysed by GC-FID equipped with a fused-silica capillary column (BPX 70: 30 m length, 0.32 mm diameter and 0.25 μm film thicknesses) (SGE Analytical Science). The injector temperature was fixed at 250°C and the oven temperature program was held at 80°C for 1 min and then ramped at 10°C min^-1^ to 235°C for 1 min.

The individual fatty acid methyl ester (FAME) peaks were identified by comparing the retention time of the FAME peaks with the retention time of the injected FAME reference standards. The peak area of the chromatogram was exploited to calculate the percentage of the total fatty acid in the oil palm root.

## Results

Obtaining adequate biomolecule samples which are reproducible and of high quality, and minimising variables are always top priority in any biological research. Oil palm is the most productive oil-bearing crop and research to understand the systems biology especially relating to its fruit development and oil productivity is of pertinent interest. Generally, oil palm RNA, DNA, proteins and metabolites are extracted independently based on research goals. Oil palm, as other free-standing plants, is supported by its root system, which acts as the main source of nutrient for growth and development. In this study, oil palm root samples were used in the development of an all-in-one extraction protocol. Later, this protocol could be applied to other part oil palm tissues such as fruit mesocarp and leaf. The all-in-one extraction protocol using oil palm root samples was adopted from Weckwerth *et al*. (2004) and Valledor *et al*. (2014) with minor modification. The protocol allowed for fast and reproducible extraction of metabolites, proteins and RNA from the oil palm root samples.

### Metabolite analysis of oil palm root samples

High throughput liquid chromatography-mass spectrometry (LC-MS) technique aims at analysis and identification of small molecules involved in metabolic reactions. Here, polar phase derived from the oil palm root was subjected to LC-MS as a benchmark of wide coverage identification of metabolites. Fig. 2 shows base peak chromatogram of oil palm root polar extract generated from LC-MS analysis. The chromatogram shows good separation and coverage of metabolites. This approach has also been established for the detection of metabolic signatures in biological samples (Zhou *et al*., 2012).

**Fig. 2.**
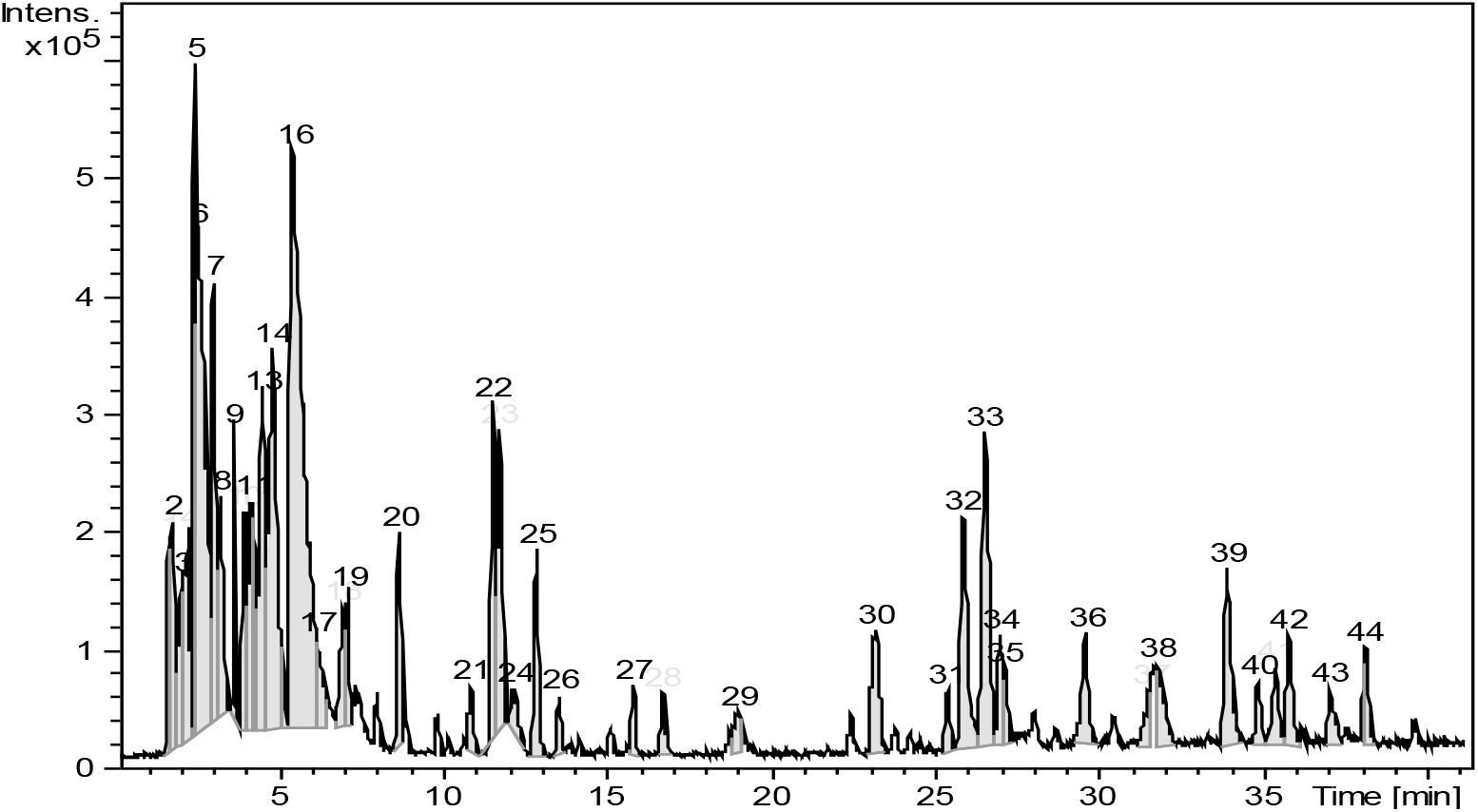
Liquid chromatography mass spectrometry (LC-MS) base peak chromatogram (BPC) of oil palm root extract in negative ionization mode. More than 40 peaks were detected in the oil palm root sample.

Further analysis was also conducted to the non-polar phase by measuring the fatty acid content in the oil palm root using gas chromatography coupled with flame ionization detector (GC-FID). Heptadecanoic acid (C17:0) was used as an internal standard. Oil palm root consists of two major fatty acids i.e. polyunsaturated and unsaturated fatty acids at 55.4% and 38.6%. The fatty acid analysis showed six major fatty acids (Fig. 3), which were palmitic acid (1.75% ± 0.04), stearic acid (55.36% ± 0.04), oleic acid (4.00% ± 0.05), linoleic acid (8.64% ± 0.08), linolenic acid (23.14% ± 0.31) and eicosanoic acid (1.08% ± 0.02) with coefficient variation (CV) percentage less than 3%, respectively. The low percentage of CV showed the mean value for the peak area for each biological replicate samples were quite close and reproducible.

**Fig. 3.**
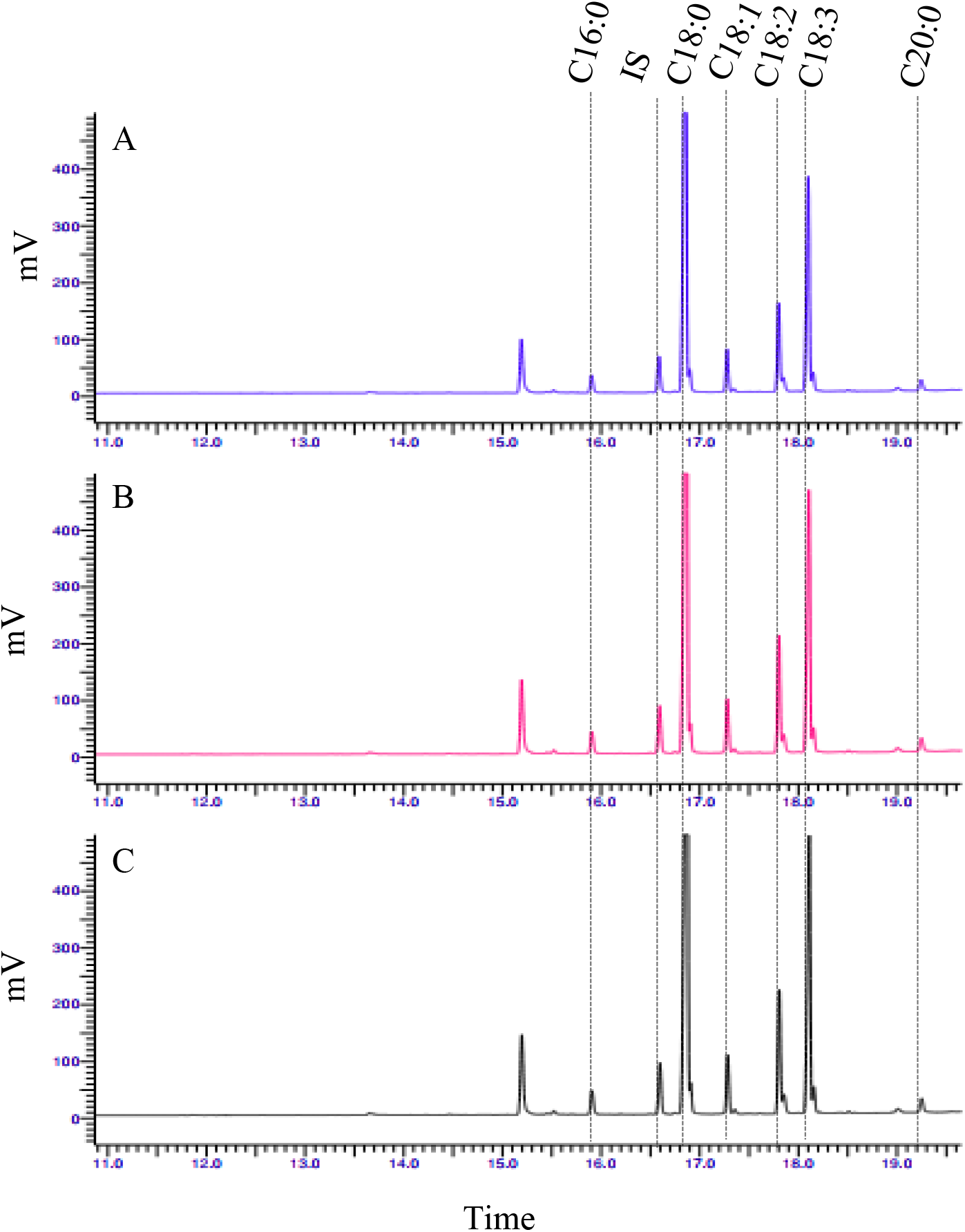
GC-FID chromatogram of fatty acid methyl esters (FAME) from the oil palm root in triplicate (Root 1, Root 2 and Root 3). Oil palm root consists of C16:0 (palmitic acid), C18:0 (stearic acid), C18:1 (oleic acid), C18:2 (linoleic acid), C18:3 (linolenic acid) and C20:0 (eicosanoic acid) fatty acids. Heptadecanoic acid (C17:0) was used as an internal standard (IS).

### Protein analysis by gel-based liquid chromatography-tandem mass spectrometry (LC-MS/MS)

The all-in-one protocol employed protein extraction buffers and phenol-based extraction to extract proteins from plant tissues as a starting material (Weckwerth *et al*., 2004; Valledor *et al*., 2014). In addition, Syahanim *et al*. (2013) has extracted proteins from oil palm roots using phenol-based extraction method. Thus, the protein extraction buffer formulation was used based on these three papers and designated as protein extraction buffer A, B and C, respectively. Accordingly, the protein extracts obtained from protein extraction buffer A, B and C were quantified using Qubit Protein Assay Kit. Referring to BSA (5 mg ml^-1^) as reference protein, a standard curve was plotted to calculate the protein concentration. Protein recoveries of 17.32 ± 1.21 μg mg^-1^ (Buffer A), 29.28 ± 2.32 μg mg^-1^ (Buffer B) and 27.40 ± 2.19 μg mg^-1^ (Buffer C) from the oil palm root were recorded, respectively (*Table 1*). Protein extraction buffer based on Weckwerth *et al*. (2004) (Buffer B) demonstrated highest protein recovery of 29.28 ± 2.32 μg mg^-1^ fresh weight (FW) tissue with a CV of 7.92% followed by protein extraction buffer based on Syahanim *et al*. (2013) with slight modification revealed a protein recovery of 27.40 ± 2.19 μg mg^-1^ FW tissue with a CV of 7.99%. The quality of extracted proteins was evaluated by sodium dodecyl sulphate-polyacrylamide gel electrophoresis (SDS-PAGE), followed by Coomassie Brilliant Blue staining (Vorreiter *et al*., 2016). Fig. 4 shows the oil palm root protein separation in one dimensional SDS-PAGE (12%). Visual comparison of protein bands in SDS-PAGE lanes corresponding to oil palm root protein extracts derived from protein extraction buffer A, B and C demonstrated distinct similarities. Gel-liquid chromatography fractionation coupled to mass spectrometry was performed on proteins extracted from buffer C in three biological replicates. Tandem mass spectrometry demonstrated a total of 90,182 MS spectrum was generated from the tryptic peptide of the oil palm root. A peptide spectrum matching was conducted against oil palm databases collection derived from in house datasets and public repository and revealed a total 1852 ± 5 identified proteins in triplicate. All peptides were validated using the percolator^®^ algorithm, based on q-value less than 1% false discovery rate (FDR). Multi-scatter plot with Pearson correlation coefficient were above 0.95, demonstrating a positive correlation in between three biological replicates (Fig. 5). Protein classification was performed via Blast2Go, Go version: Jan 1 2020 and the identified oil palm root protein datasets were classified in three groups of biological process, cellular component and molecular function (Fig. 6). Digested proteins derived from simultaneous extraction were subjected to high resolution accurate mass (HRAM) mass spectrometry as demonstrated by Vorreiter *et al*. (2016) and Weckwerth *et al*. (2004) and revealed 1091 and 297 identified proteins from the human T-cell lines and *Arabidopsis* sample respectively.

**Fig. 4.**
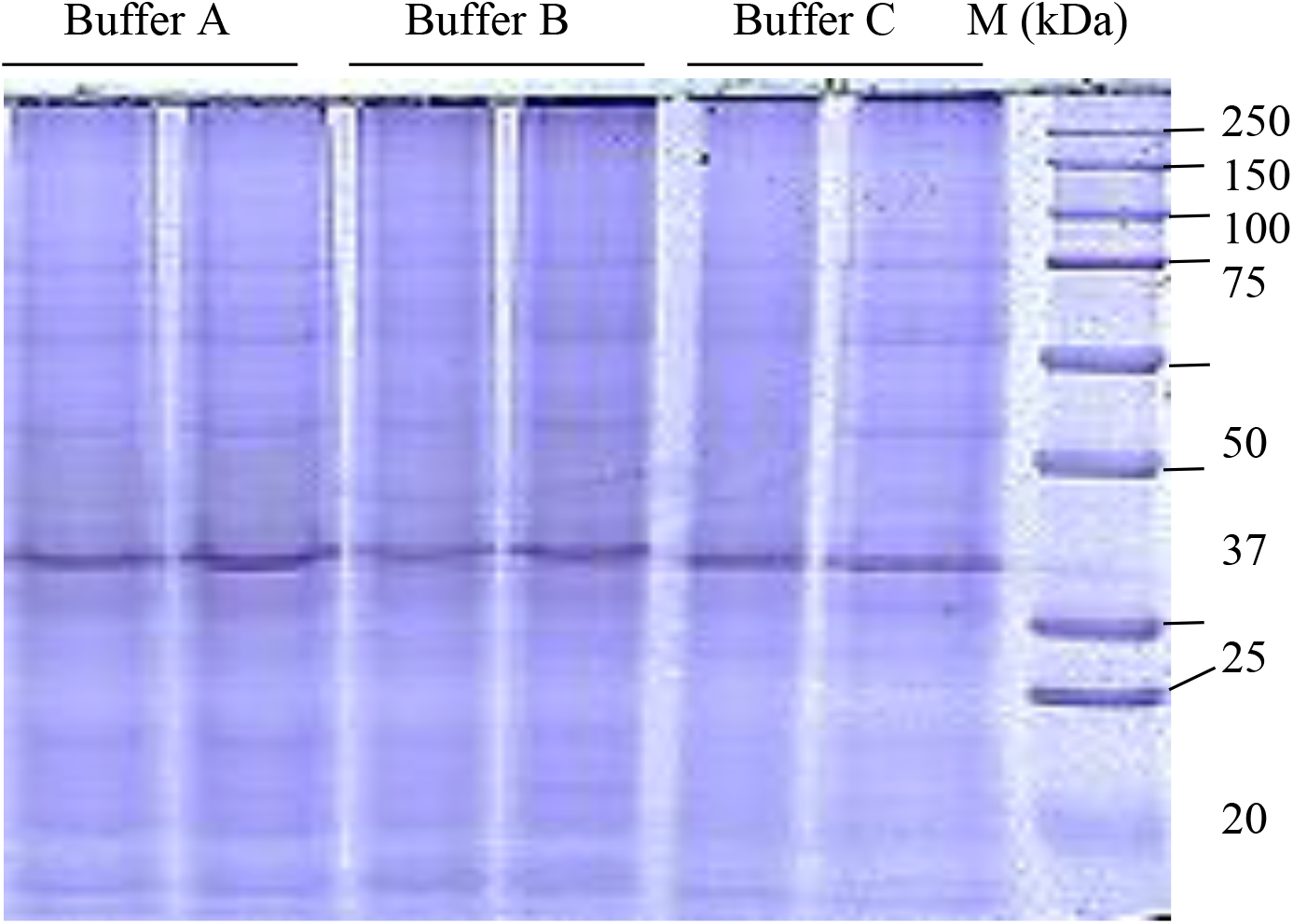
One dimensional gel electrophoresis of the oil palm root samples (25 μg) extracted using protein extraction buffer A, B, and C, respectively. M: Kaleidoscope protein marker.

**Fig. 5.**
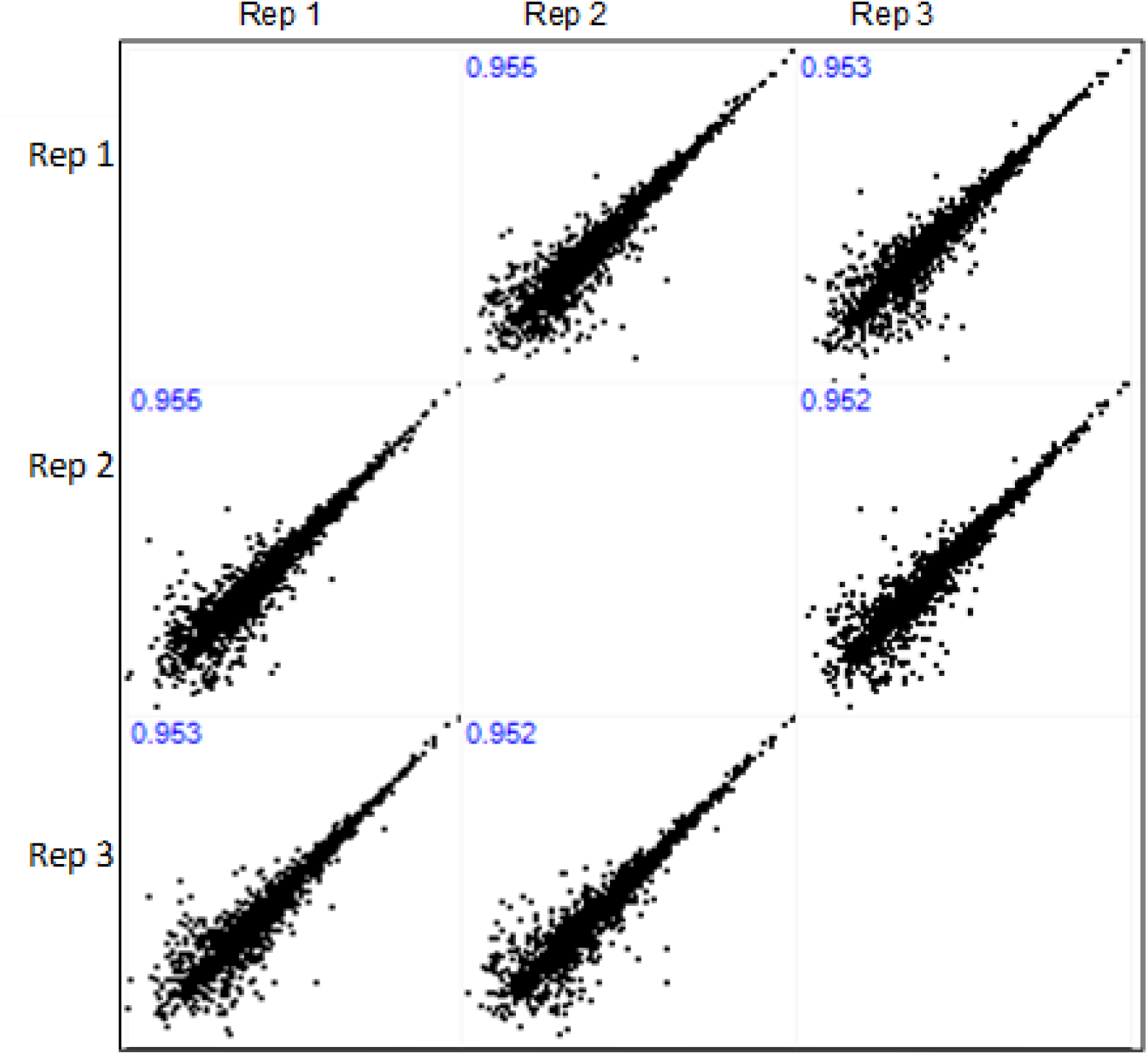
Multi-scatter plots with Pearson correlation values of above 0.95, suggesting a good correlation between data obtained from biological replicates samples of oil palm root.

**Fig. 6.**
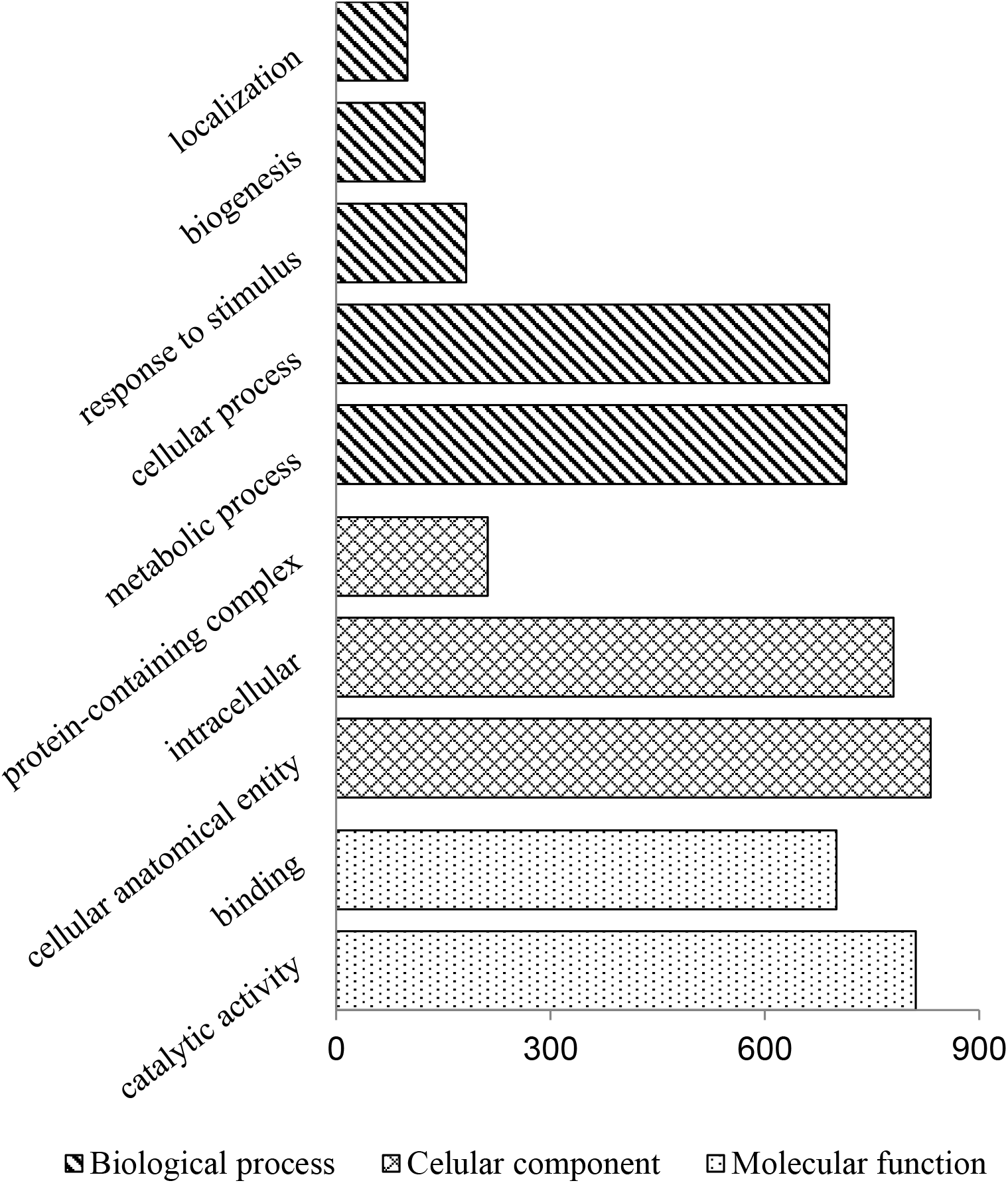
Annotation based on Gene Ontology terms using Blast2Go of amino acid sequences derived from the oil palm root proteome.

**Table 1.**
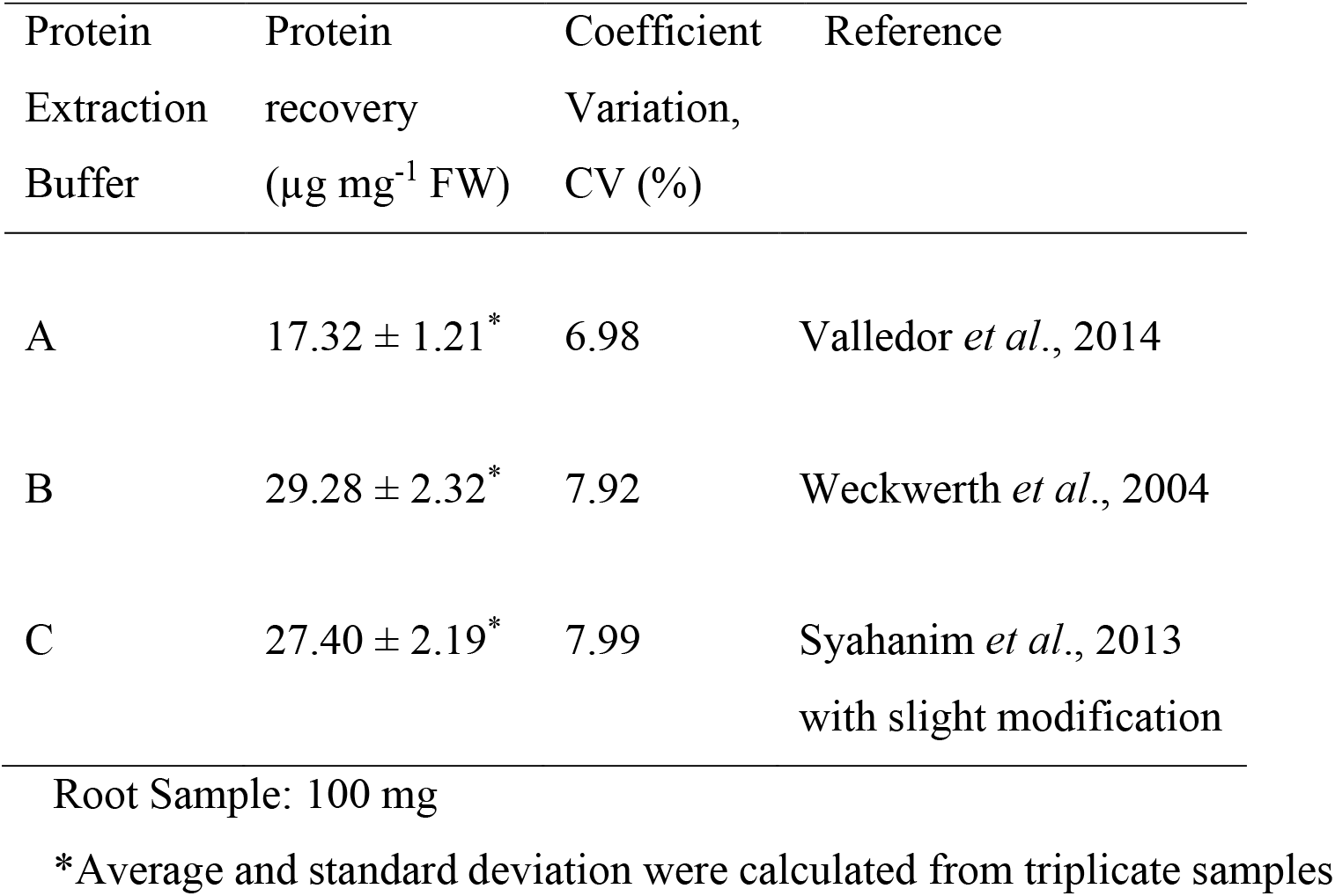
Oil palm root protein recovery extracted using three different protein extraction buffers.

### RNA quality assessment

RNA was extracted twice from the buffer phase by using chloroform. The quality of RNA derived from oil palm root was assessed using a NanoDrop One system (Thermo Scientific, Wilmington, DE, USA) and the purity was estimated by the A260:A280 ratio. The RNA recovery was 180.25 ± 8.95 ng mg^-1^ (n=3) and demonstrated a purity of 1.9-2.0 based on the ratio of A260:A280. A good RNA integrity number value is required for downstream applications such as sequencing and real time PCR. RIN values range from 1.0 (degraded) to 9.0 (intact). RIN value is calculated and generated from Agilent 2100 Bioanalyzer. The integrity of RNA extracted from the oil palm root using sequential protocol generated a RIN value of 6.7 (Fig. 7). Therefore, our results suggest that the all-in-one extraction method employed in this study is capable of producing good quality of RNA from oil palm root tissue adequate for downstream analysis.

**Fig. 7.**
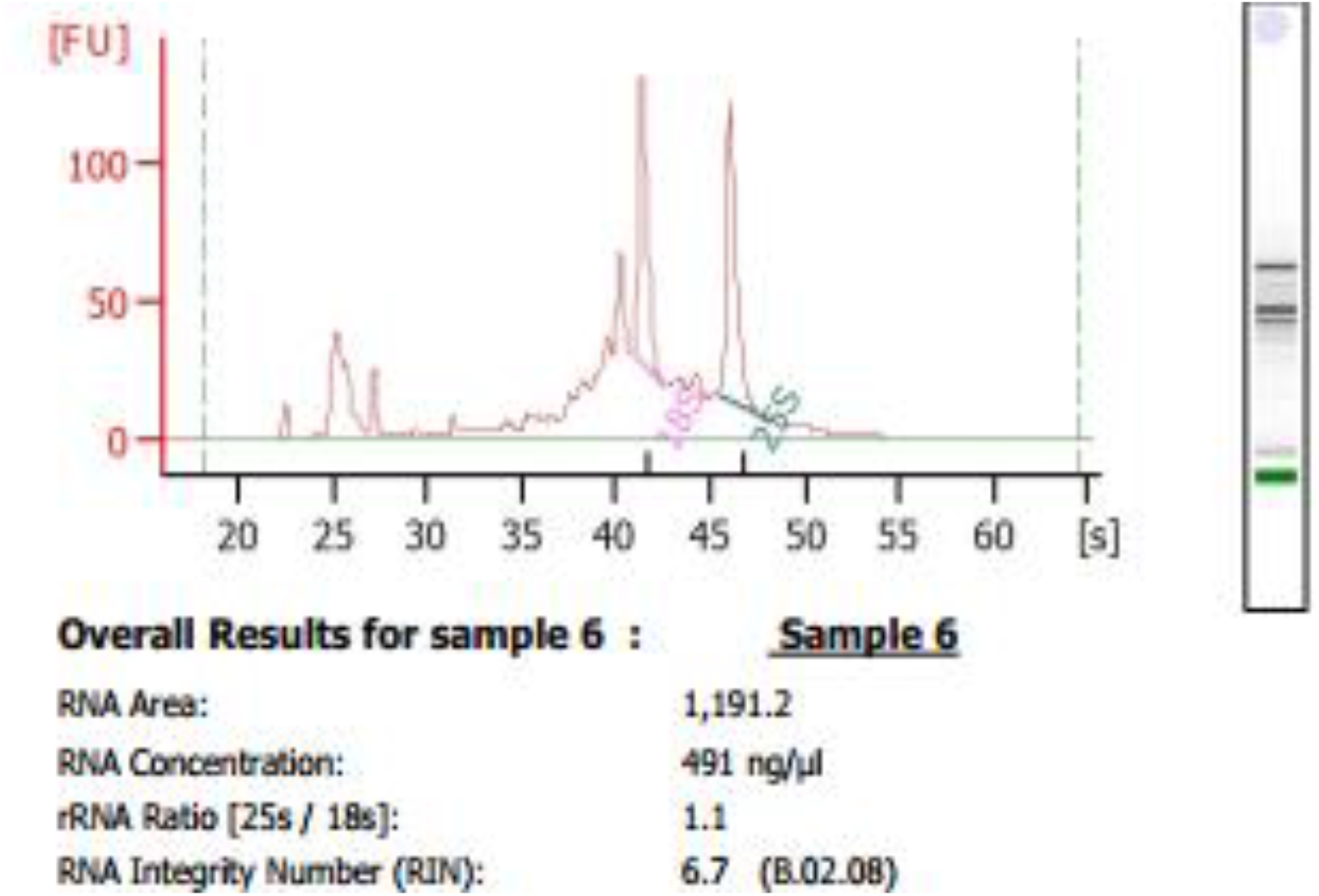
A representative image of electropherogram of total RNA sample from the oil palm root evaluated using the Bioanalyzer (Agilent Technologies Inc).

## Discussion

Several reports have demonstrated the compatibility of biomolecule components extracted using simultaneous extraction protocols for various multi-omics approaches. Chomczynski (1993) invented a mono-phase solution which consisted of acid guanidine thiocyanate-phenol-chloroform for the single-step simultaneous isolation of RNA, DNA and proteins from animal tissues. Electrophoretic pattern of RNA showed two distinct ribosomal bands, as an indicator for authenticity of the total RNA. A column-based commercial kit was also tested for co-extraction of RNA and proteins as described by Morse *et al*. (2006). The team used RNeasy^®^ Mini Kit (QIAGEN) to extract RNA from cultured cell lines. After column centrifugation, the flow-through was retained and subjected to protein precipitation. By performing this procedure, concurrent extraction of RNA and proteins from the experimental sample was efficiently conducted. Method optimisation for simultaneous extraction of RNA, DNA, proteins and metabolites was reported using human T-cells and hepatocytes cell lines in which three methods were tested (Vorreiter *et al*., 2016). The first method was based on Weckwerth *et al*. (2004). For comparison, a second method was tested using commercial buffer, TRI REAGENT™ (www.sigmaaldrich.com) according to the manufacturer’s protocol. The third method employed phenol/chloroform for RNA, DNA and proteins with an additional step included to extract metabolites using methanol/chloroform. The suitability of the extraction methods for the biomolecules was evaluated and it was observed that the third method adequately produced good quality of DNA, RNA, proteins and metabolites from the cell lines. Weckwerth *et al*. (2004) and Morgenthal *et al*. (2005) have demonstrated a novel extraction protocol for integrated extraction of biomolecules from *Arabidopsis thaliana* leaves. Valledor *et al*. (2014) used several plants to develop an improved universal protocol for isolation of biomolecule component in a single sample of *Chlamydomonas*, *Arabidopsis*, *Populus* and *Sinus*. Additionally, metabolite extraction was conducted based on Weckwerth *et al*. (2004) while subsequent isolation of proteins and RNA was performed using commercial kits.

Despite recent advancement in oil palm proteomics and metabolomics research (Ramli *et al*., 2016), multi-omics analysis of the same sample of oil palm is still challenging. In this report, we describe a novel protocol for the sequential extraction of metabolites, proteins and RNA from the same undivided oil palm root sample to facilitate multi-omics analysis and meaningful data integration. Simultaneous extraction of metabolites, proteins and RNA from the oil palm tissue was exploited using the established protocol by Weckwerth *et al*. (2004) and Valledor *et al*. (2014) with slight modification during RNA and proteins extraction. To isolate RNA from the oil palm samples, a classical phenol/chloroform protocol (Weckwerth *et al*., 2004) is preferred than using a silica-based column (Valledor *et al*., 2014) due to the rising cost of commercially available columns. However, the classical RNA extraction is prone to contamination of phenol and salts that may affect the RNA quality for downstream application. To overcome this, we introduced a second chloroform washing step, as well as a twice washing step as described by Toni *et al*. (2016). For animal and human tissues, RIN value of 9.0 is needed for further analysis. For plants, RIN value of more than 5 is acceptable for further downstream analysis (Fleige and Pfaffl, 2006). This is due to the fact that plant species have diverse ribosomal RNA sizes (5S, 8S, 16S, 23S and 25S) from mitochondria, cytosol and chloroplast which contribute to the complexity in RIN value. The classical RNA extraction described in this paper produced adequate RNA of acceptable quality from oil palm root tissue for downstream analysis.

In the following sections, we provide an exemplary of the multi-level extraction of proteins from oil palm root using different buffers as previously reported by other groups (Weckwerth *et al*., 2004; Syahanim *et al*., 2013; Valledor *et al*., 2014). However, the performance of the protein extraction buffer for the integrated extraction protocol has never been reported for an oil palm tissue. In the present study, after removing the solvent phase which contained metabolites, the remaining solid pellet was used for protein extraction. This step employed extraction buffers and phenol to extract proteins as described previously for plant tissues (Weckwerth *et al*., 2004; Valledor *et al*., 2014). Syahanim *et al*. (2013) also reported an independent phenol-based extraction method to extract proteins from oil palm roots. The protein extraction buffer formulation described in the above three papers involved the usage of sucrose as a phase conversion and reducing agent. The protein extraction buffer formulation reported by Weckwerth *et al*. (2004) produced the highest protein recovery. The buffer formulation reported by Syahanim *et al*. (2013) was also included in this work, although it produced slightly lower protein recovery. The quality of extracted proteins was evaluated by SDS-PAGE, followed by Coomassie Brilliant Blue staining and revealed the same protein bands during the SDS-PAGE separation. Nonetheless, the results indicated that the three protein extraction buffers can be used for protein extraction using oil palm root samples.

Thus, the prospect of extracting metabolites, proteins and RNA from oil palm tissues using an all-in-one extraction protocol reported in this paper is very attractive for achieving fast and high quality biomolecule extracts, in a single experiment. In systems biology studies, the data integration derived from multi-omics analysis (genomics, transcriptomics, proteomics and metabolomics) has been shown to provide more complete and informative view of biological pathways. Thus, the developed integration protocol could be explored further in other oil palm tissues such as mesocarp and leaf. This will enable comprehensive analysis using various omics platforms to fill knowledge gaps in oil palm systems biology.

## Abbreviations

RNA: Ribonucleic acid
LC-MS/MS: liquid chromatographytandem mass spectrometry
GC-FID: gas chromatography-flame ionization detector
SDS-PAGE: sodium dodecyl sulphate-polyacrylamide gel electrophoresis
FAME: fatty acid methyl esters

## Acknowledgement

The authors would like to thank the Director-General of the Malaysian Palm Oil Board (MPOB) for permission to publish this paper. We thank Dr Mohd Din Amiruddin of Breeding and Genetics Unit, Advanced Biotechnology and Breeding Centre (ABBC) MPOB for assistance in providing the oil palm root samples. We also would like to thank Asmahani Azira of Malaysian Genome Institute for the technical assistance with Orbitrap Fusion mass spectrometry. We acknowledge the technical support of Genomics Unit, Bioinformatics Unit and Proteomics and Metabolomics Unit of ABBC throughout this study. We would like to express our gratitude to Dr Ravigadevi Sambantahmurthi for her valuable comments.

